# The polypharmacological profiles of xanomeline and N-desmethylxanomeline

**DOI:** 10.1101/2025.11.20.689572

**Authors:** Kensuke Sakamoto, Xi-Ping Huang, Talia L. Albert, Paloma M. Knobloch, Grace K. Foland, Bryan L Roth

**Affiliations:** NIMH Psychoactive Drug Screening Program and the Department of Pharmacology, UNC Chapel Hill Medical School, Chapel Hill, NC, USA 27705

## Abstract

The muscarinic agonist xanomeline in combination with the peripherally restricted muscarinic antagonist trospium has recently been approved for treatment of schizophrenia. Xanomeline represents the first approved antipsychotic drug without apparent activity at D2-dopamine receptors. In humans, xanomeline is reported to be metabolized to N-desmethylxanomeline, which has a similar pharmacokinetic profile to xanomeline, although its pharmacology has not been reported. We discovered that xanomeline and N-desmethylxanomeline have potent agonist and antagonist actions at many biogenic amine G protein coupled receptors. These results suggest that at least some of the actions of xanomeline and N-desmethylxanomeline could be mediated by off-target actions at serotonergic, dopaminergic, histaminergic, adrenergic and other receptors. We discuss the potential implications of these findings.

## Introduction

Xanomeline was first developed as a muscarinic agonist for Alzheimer’s Disease ^1^ but because of side-effects due to activation of peripheral muscarinic receptors was not advanced. Parenthetically, xanomeline was reported to improve both cognitive and psychotic symptoms in both Alzheimer’s Disease ^2^ and schizophrenia ^3^. Ultimately, xanomeline was combined with trospium—a peripheral muscarinic antagonist—and demonstrated to improve symptoms of schizophrenia with few side-effects due mainly to peripheral muscarinic agonism ^4,5^. This combination of xanomeline and trospium was recently approved by the US FDA for treating schizophrenia ^6^. Xanomeline thus represents a breakthrough as the first non-D2 dopamine receptor modulator approved to treat schizophrenia. Given the lack of significant reported D2 antagonist activity, xanomeline has minimal motoric side-effects and is weight neutral when compared with other approved antipsychotic drugs ^4,5^.

We have been interested in the potential polypharmacology of CNS-active compounds for many years ^7^ and noted that xanomeline was reported to have activities at several other molecular targets which have been implicated in antipsychotic drug actions ^8^. Xanomeline is also known to be metabolized to N-desmethylxanomeline which was reported to achieve peak concentrations similar to the parent drug xanomeline in human volunteers ^9^. To our knowledge, there are no published data on the pharmacological profile of N-desmethylxanomeline although one report from 1995 states it is a “pharmacologically active metabolite” ^9^.

Accordingly, we carried out a comprehensive off-target analysis of xanomeline and N-desmethylxanomeline via radioligand binding and functional assays at many G protein coupled receptors (GPCRs) and other CNS targets. We discovered that both compounds have rich polypharmacological profiles which could be important for their actions in humans.

## Methods

### Compounds

Xanomeline and N-desmethyl xanomeline were purchased from MedChemExpress (HY-105182) and EnamineStore (Z4444421058), respectively.

### PRESTO-Tango GPCRome Screen and Follow-up Assays

Off-target agonism across the human GPCRome was profiled using PRESTO-Tango, following established protocols with minor adaptations ^10–12^. HTLA cells were seeded at 10,000 cells per well in 40 µL Dulbecco’s Modified Eagle Medium (DMEM) supplemented with 1% dialyzed fetal bovine serum (dFBS) in poly-L-lysine-coated, white, clear-bottom 384-well plates. After ~6 h recovery, cells were transfected with 20 ng plasmid DNA per well and incubated overnight. The next day, 10 µL of test compound (prepared in DMEM + 1% dFBS) was added, and plates were incubated overnight. Media and compounds were removed, and 20 µL per well of Bright-Glo luciferase reagent diluted in assay buffer (Hank’s Balanced Salt Solution (HBSS), 20 mM HEPES, pH 7.4) was added. After 20 min at room temperature (dark), luminescence was recorded.

Each plate included the dopamine D2 receptor (DRD2) stimulated with 100 nM quinpirole as a positive control. For the GPCRome screen, each receptor was tested in quadruplicate at 0 and 10 µM compound (final). Responses were expressed as fold change relative to the basal (0 µM) condition. Follow-up concentration–response assays were performed under identical conditions except that serial dilutions of compounds (maximum ~30 µM, final) were added to generate full concentration–response curves.

### Radioligand Binding Assays

Competitive radioligand binding assays were performed at the NIMH Psychoactive Drug Screening Program (PDSP) using membranes from transiently transfected HEK293 cells in 96-well format as described ^13^. Complete, target-specific protocols are available at the PDSP website (https://pdsp.unc.edu/pdspweb/?site=assays). Primary screenings were run at a single test concentration (10 µM) in quadruplicate. Compounds producing ≥50% inhibition at 10 µM were advanced to secondary assays to determine equilibrium binding affinity. Secondary assays were conducted in triplicate using 12-point serial dilutions (0, 0.1, 0.3, 1, 3, 10, 30, 100, 300 nM; 1, 3, 10 µM). Assays were performed in a final volume of 125 µL per well in binding buffer, as detailed in the NIMH PDSP protocols (e.g., standard binding buffer: 50 mM Tris-HCl, pH 7.4, 10 mM MgCl_2_, 0.1 mM EDTA). Radioligand concentrations were chosen near the target-specific K_d_. Total and nonspecific binding were defined in the absence and presence, respectively, of 10 µM reference ligand specific for each receptor. After 90 min incubation at room temperature, reactions were terminated by vacuum filtration onto 0.3% polyethyleneimine-pretreated UniFilter-96 GF/C Microplates using a FilterMate harvester, washed three times with ice-cold buffer (50 mM Tris HCl, pH 7.4), dried, treated with MicroScint-O liquid scintillation cocktails, and counted on a MicroBeta counter. Equilibrium inhibition constants (K_i_) were calculated from fitted IC_50_ values using the Cheng–Prusoff equation ^14^ with the assay-specific radioligand concentration and K_d_.

### G-Protein Dissociation (BRET2)

BRET2 assays followed prior methods ^15,16^ with minor adjustments. HEK293 cells were maintained in DMEM containing 10% FBS, 100 U mL^−1^ penicillin, and 100 µg mL^−1^ streptomycin at 37 °C in 5% CO_2_. For each experiment, 2-3 × 10^6^ cells were plated per 6-cm dish and transfected 24 h later using TransIT-2020 (Mirus; 3 µL per µg DNA). Plasmids encoding the receptor, the appropriate Gα-RLuc8, Gβ, and Gγ-GFP2 (as in the cited studies) were co-transfected at 250 ng each per dish. After 24 h, cells were detached with 0.05% trypsin-EDTA, resuspended in DMEM with 1% dFBS, and seeded at 1 × 10^4^ cells per well into poly-L-lysine-coated white 384-well plates. Twenty-four hours later, plates were backed with white film, media were removed, and wells were washed once with 20 µL assay buffer (HBSS, 20 mM HEPES, pH 7.4, 0.1% BSA). Serial dilutions of ligands (top concentration up to 30 µM, final) were prepared in assay buffer. To each well, 20 µL buffer containing 5 µM coelenterazine-400a and 10 µL ligand solution were added. After 15 min at 37 °C, emission at 395 nm (donor) and 510 nm (acceptor) was recorded on a PHERAstar FSX plate reader. BRET2 ratios (GFP2/RLuc8) were normalized to plate controls; concentration–response data were fit in GraphPad Prism v10 using a three-parameter logistic model to derive EC_50_ and Emax. To estimate the transduction coefficient log(τ/KA), we additionally fit the Black-Leff operational model^17^. For antagonist-mode experiments, cells were preincubated with test compounds and challenged with an agonist at the receptor-specific EC_80_ (e.g., histamine for histamine receptors) 15 min before reading.

### β-Arrestin-2 Recruitment (BRET1)

BRET1 assays followed prior methods ^18^. Receptors were fused C-terminally to RLuc8 and co-transfected with GRK2 and mVenus-β-arrestin-2 at a 1:1:5 plasmid ratio. Assay conditions and readings matched BRET2, except that coelenterazine-h was used as the luciferase substrate. Emission at 475 nm (donor) and 535 nm (acceptor) was recorded on a PHERAstar FSX plate reader.

### GloSensor cAMP Assay

Intracellular cAMP regulations downstream of Gαs or Gαi were quantified using the GloSensor-22F reporter (Promega) as described ^13^. HEK293 cells were transiently co-transfected with GloSensor and the indicated receptor constructs, then seeded the next day at 1.0 × 10^4^ cells per well in poly-L-lysine-coated white, clear-bottom 384-well plates (40 µL DMEM + 1% dFBS). After 24 h, media were replaced with 20 µL per well of assay buffer (HBSS, 20 mM HEPES, pH 7.4, 0.1% BSA) containing serially diluted test ligands and 3 mM D-luciferin (GoldBio). Plates were incubated for 15 min at room temperature, and basal luminescence was read on a SpectraMax L luminometer to assess Gαs-mediated cAMP elevation. To assess Gαi-mediated cAMP inhibition, isoproterenol (10 µL per well; 100 nM final) was added to activate endogenous β_2_-adrenergic receptors; luminescence was recorded after an additional 15 min. For antagonist-mode experiments, cells were preincubated with test compounds and then challenged with an agonist at its EC_80_ (e.g., quinpirole for dopamine receptors). Data were normalized to plate controls and fit in GraphPad Prism v10 using a three-parameter logistic model.

### Calcium Mobilization Assay

Calcium flux was measured with Fluo-4 Direct (Invitrogen) as described ^19^. For 5-HT_2_A/5-HT_2_B/5-HT_2_C, HEK293 cell lines stably expressing the corresponding receptor were maintained in DMEM with 10% FBS, 100 U mL^−1^ penicillin, 100 µg mL^−1^ streptomycin, 100 µg mL^−1^ hygromycin B, and 15 µg mL^−1^ blasticidin at 37 °C/5% CO_2_. Receptor expression was induced with tetracycline (1.5 µg mL^−1^), and cells were seeded into 384-well black plates at ~10,000 cells per well in DMEM + 1% dFBS with antibiotics. For CHRM1/CHRM3/CHRM5, HEK293 cells were transiently transfected with the respective receptor constructs under comparable conditions.

Twenty-four hours later, cells were loaded with 20 µL per well of Fluo-4 Direct supplemented with 2 mM probenecid in drug buffer (HBSS, 20 mM HEPES, 0.1% BSA, 0.01% ascorbic acid, pH 7.4) for 1 h at 37 °C, followed by 10 min at room temperature in the dark. Ligands were prepared at 3× in drug buffer (top concentration ~30 µM). Baseline fluorescence was acquired for 10 s (1 read s^−1^) on a FLIPR-Tetra, followed by automated addition of 10 µL ligand (3×) and continued acquisition for 120 s (1 read s^−1^). Maximal fluorescence over 120 s was converted to fold change relative to baseline (mean of the initial 10 reads). Concentration–response curves were normalized to controls and fit in GraphPad Prism v10 using a three-parameter logistic model.

## Results

### Radioligand binding assays reveal multiple molecular targets for xanomeline and N-desmethylxanomeline

In initial experiments, we performed a screen of xanomeline and N-desmethylxanomeline at 45 distinct molecular targets via radioligand binding assays as previously detailed ^13^ (Fig 1; Table 1) using the resources of the National Institute of Mental Health Psychoactive Drug Screening Program (PDSP). Table 1 also shows our results with those and also those values previously reported by Murphy et al ^9^.

**FIGURE 1.**
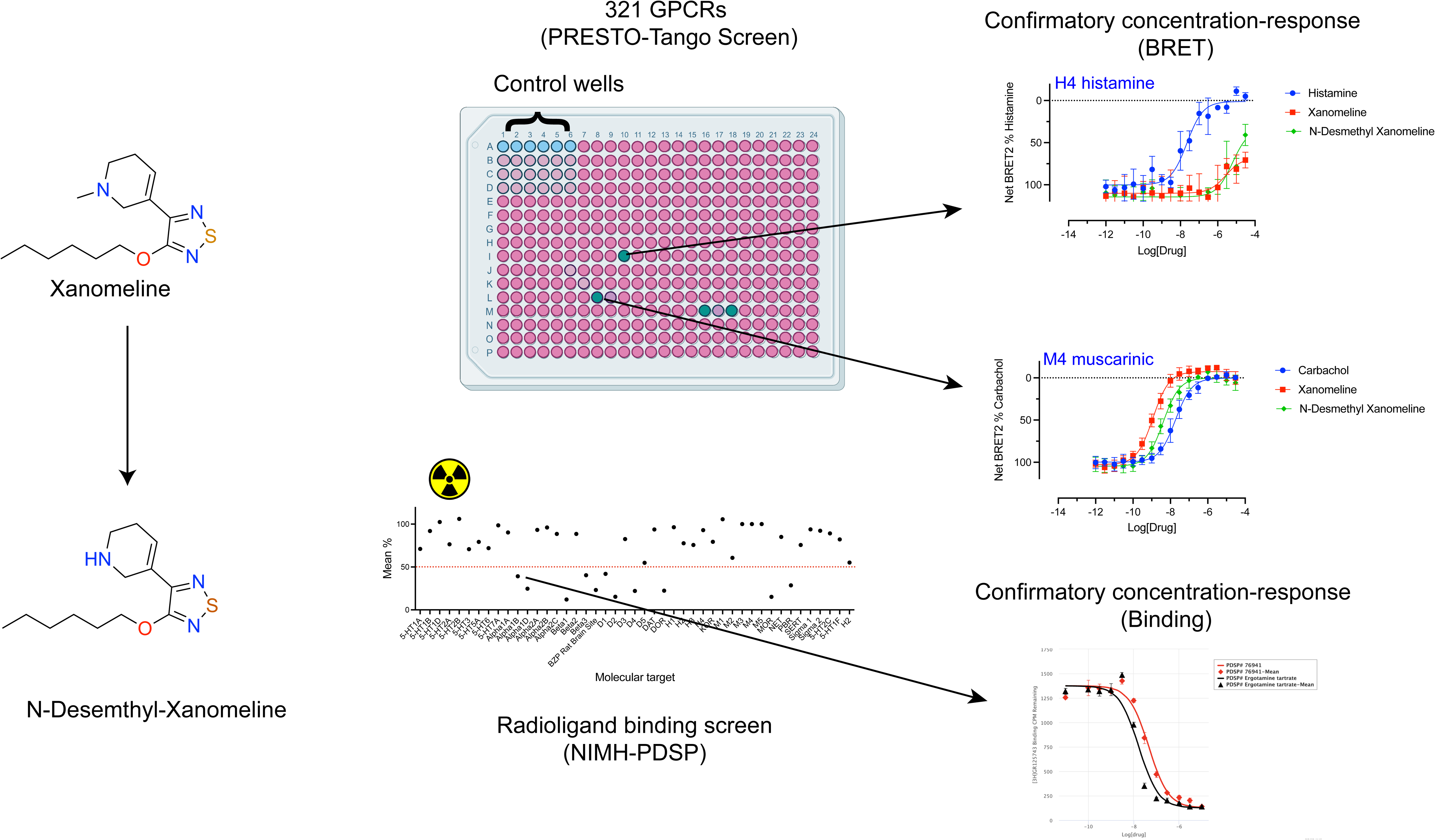
Strategy for identifying potential molecular targets for Xanomeline and N-desmethyl-Xanomeline. As shown, Xanomeline and N-desemethyl-Xanomeline were profiled using the PRESTO-TANGO and NIMH-PDSP profiling resources. Where activities were >50% in the initial NIMH-PDSP radioligand binding screen at a final concentration of 10 μM were detected, follow-up studies quantified apparent Ki values by radioligand binding assays. Where activities were > 2-fold I the PRESTO-TANGO screen, follow-up dose-response studies in BRET assays were used to confirm activities.

**TABLE 1.**
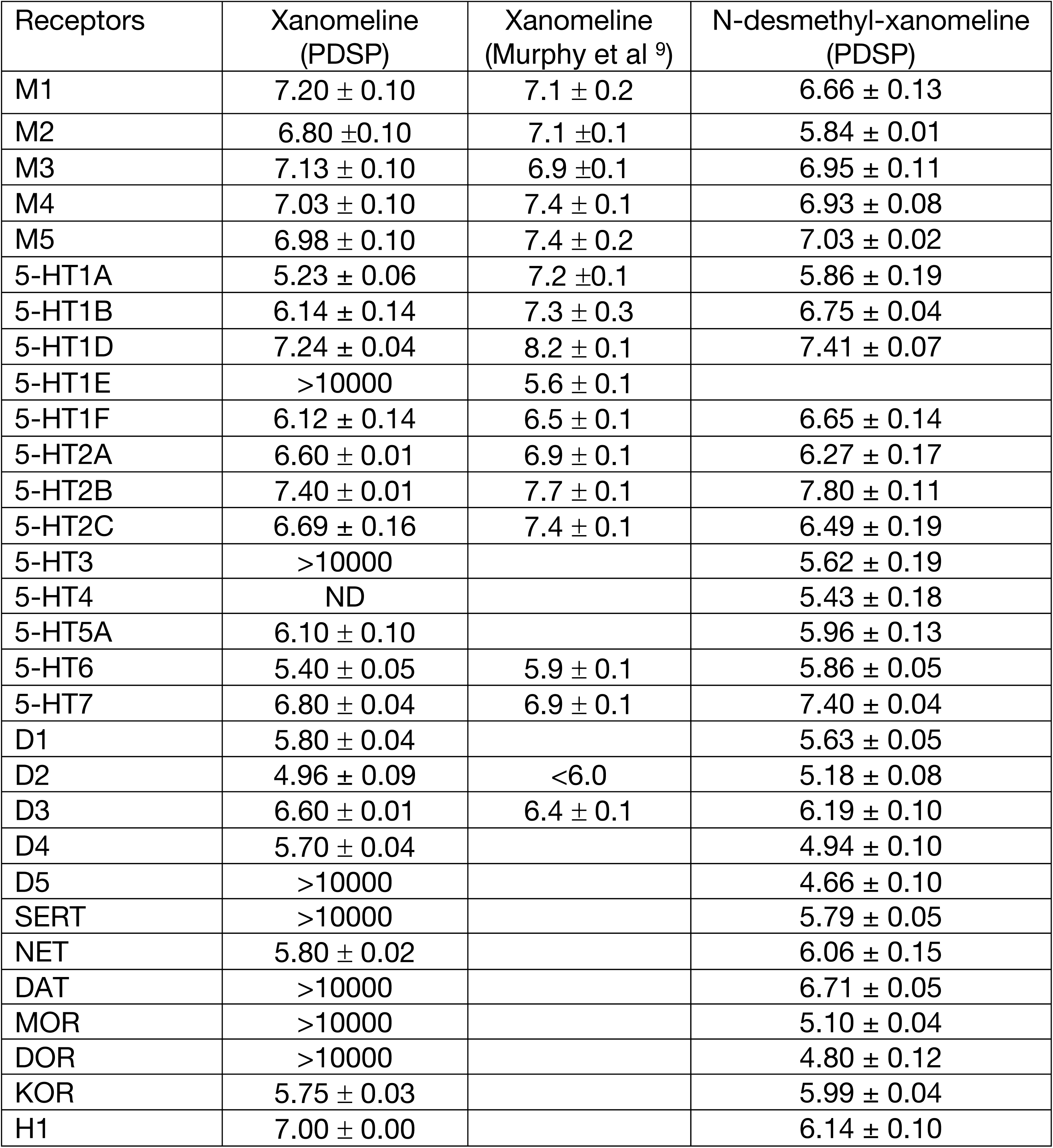
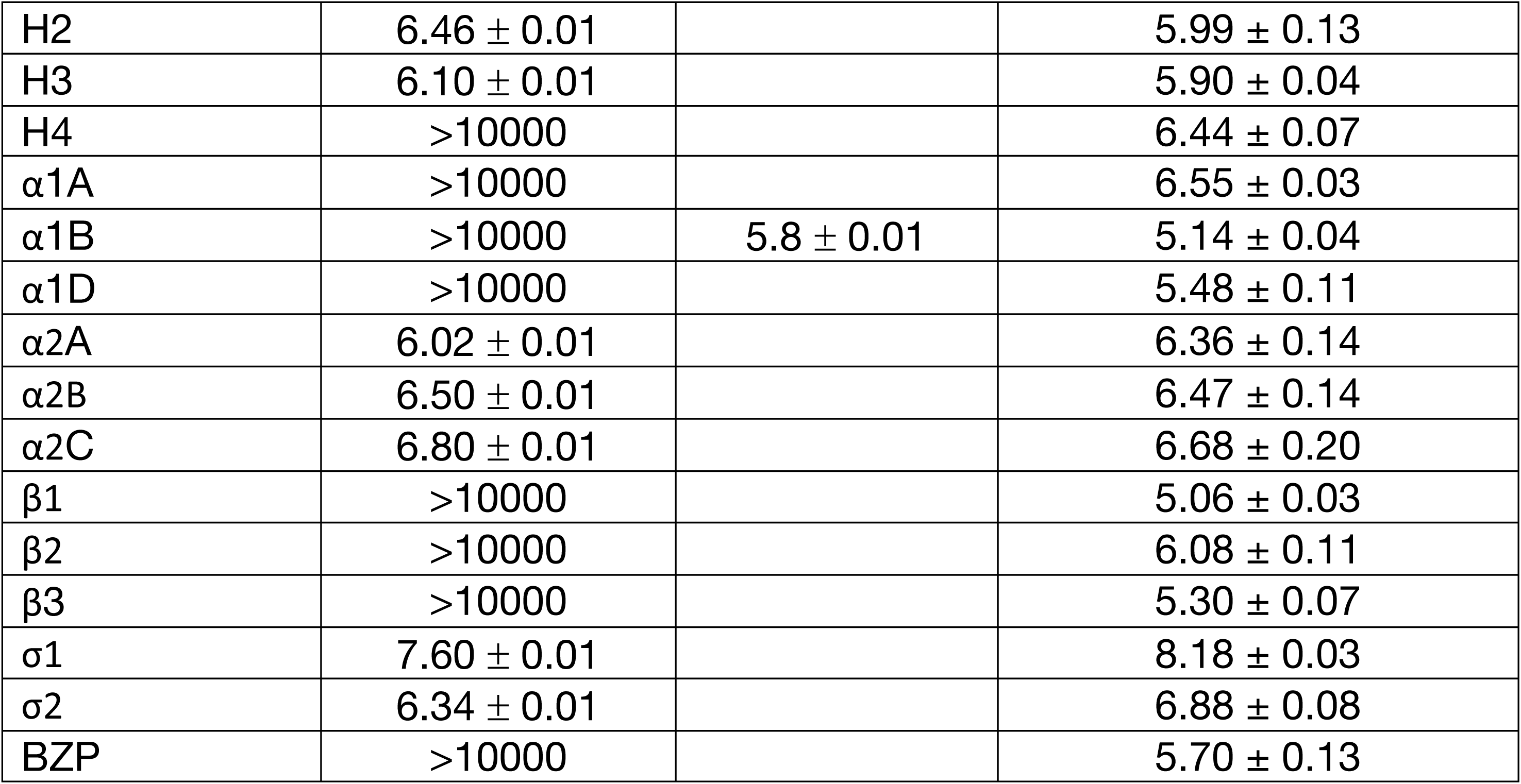
BINDING AFFINITIES OF XANOMELINE AND N-DESMETHYL-XANOMELINE AT COMMON CNS TARGETS. Xanomeline and its metabolite N-desmethyl xanomeline binding affinity (pK_i_) are reported. Published xanomeline binding affinity from Murphy et al ^9^ were included for comparison. Results represented mean ± SEM from a minimum of 3 independent assays, each in triplicate. ND=not determined.

| Receptors | Xanomeline (PDSP) | Xanomeline (Murphy et al <sup>9</sup>) | N-desmethyl-xanomeline (PDSP) |
| --- | --- | --- | --- |
| M1 | 7.20 ± 0.10 | 7.1 ± 0.2 | 6.66 ± 0.13 |
| M2 | 6.80 ± 0.10 | 7.1 ± 0.1 | 5.84 ± 0.01 |
| M3 | 7.13 ± 0.10 | 6.9 ± 0.1 | 6.95 ± 0.11 |
| M4 | 7.03 ± 0.10 | 7.4 ± 0.1 | 6.93 ± 0.08 |
| M5 | 6.98 ± 0.10 | 7.4 ± 0.2 | 7.03 ± 0.02 |
| 5-HT1A | 5.23 ± 0.06 | 7.2 ± 0.1 | 5.86 ± 0.19 |
| 5-HT1B | 6.14 ± 0.14 | 7.3 ± 0.3 | 6.75 ± 0.04 |
| 5-HT1D | 7.24 ± 0.04 | 8.2 ± 0.1 | 7.41 ± 0.07 |
| 5-HT1E | >10000 | 5.6 ± 0.1 |  |
| 5-HT1F | 6.12 ± 0.14 | 6.5 ± 0.1 | 6.65 ± 0.14 |
| 5-HT2A | 6.60 ± 0.01 | 6.9 ± 0.1 | 6.27 ± 0.17 |
| 5-HT2B | 7.40 ± 0.01 | 7.7 ± 0.1 | 7.80 ± 0.11 |
| 5-HT2C | 6.69 ± 0.16 | 7.4 ± 0.1 | 6.49 ± 0.19 |
| 5-HT3 | >10000 |  | 5.62 ± 0.19 |
| 5-HT4 | ND |  | 5.43 ± 0.18 |
| 5-HT5A | 6.10 ± 0.10 |  | 5.96 ± 0.13 |
| 5-HT6 | 5.40 ± 0.05 | 5.9 ± 0.1 | 5.86 ± 0.05 |
| 5-HT7 | 6.80 ± 0.04 | 6.9 ± 0.1 | 7.40 ± 0.04 |
| D1 | 5.80 ± 0.04 |  | 5.63 ± 0.05 |
| D2 | 4.96 ± 0.09 | <6.0 | 5.18 ± 0.08 |
| D3 | 6.60 ± 0.01 | 6.4 ± 0.1 | 6.19 ± 0.10 |
| D4 | 5.70 ± 0.04 |  | 4.94 ± 0.10 |
| D5 | >10000 |  | 4.66 ± 0.10 |
| SERT | >10000 |  | 5.79 ± 0.05 |
| NET | 5.80 ± 0.02 |  | 6.06 ± 0.15 |
| DAT | >10000 |  | 6.71 ± 0.05 |
| MOR | >10000 |  | 5.10 ± 0.04 |
| DOR | >10000 |  | 4.80 ± 0.12 |
| KOR | 5.75 ± 0.03 |  | 5.99 ± 0.04 |
| H1 | 7.00 ± 0.00 |  | 6.14 ± 0.10 |
| H2 | $6.46 \pm 0.01$ | | $5.99 \pm 0.13$ |
| H3 | $6.10 \pm 0.01$ | | $5.90 \pm 0.04$ |
| H4 | $>10000$ | | $6.44 \pm 0.07$ |
| $\alpha 1A$ | $>10000$ | | $6.55 \pm 0.03$ |
| $\alpha 1B$ | $>10000$ | $5.8 \pm 0.01$ | $5.14 \pm 0.04$ |
| $\alpha 1D$ | $>10000$ | | $5.48 \pm 0.11$ |
| $\alpha 2A$ | $6.02 \pm 0.01$ | | $6.36 \pm 0.14$ |
| $\alpha 2B$ | $6.50 \pm 0.01$ | | $6.47 \pm 0.14$ |
| $\alpha 2C$ | $6.80 \pm 0.01$ | | $6.68 \pm 0.20$ |
| $\beta 1$ | $>10000$ | | $5.06 \pm 0.03$ |
| $\beta 2$ | $>10000$ | | $6.08 \pm 0.11$ |
| $\beta 3$ | $>10000$ | | $5.30 \pm 0.07$ |
| $\sigma 1$ | $7.60 \pm 0.01$ | | $8.18 \pm 0.03$ |
| $\sigma 2$ | $6.34 \pm 0.01$ | | $6.88 \pm 0.08$ |
| BZP | $>10000$ | | $5.70 \pm 0.13$ |

We found that xanomeline had highest affinity for the α1 receptor (Ki=29 nM), followed by the 5-HT_2B_ serotonin receptor (Ki=40 nM) and then various muscarinic acethylcholine receptors (Ki’s range from 50-158 nM). Interestingly, appreciable affinity was also revealed for several molecular targets implicated in antipsychotic drug actions including the 5-HT_1A_ ^20^, 5-HT_2A_ ^21^ and 5-HT_7_ ^22,23^ serotonin receptors along with D3-dopamine receptors ^24^. Our results for those molecular targets interrogated by Murphy et al ^9^ were similar to values they reported (Table 1).

We next evaluated N-desmethylxanomeline at these same molecular targets as there are no published data on this compound’s pharmacology (Table 1). In general, N-desmethylxanomeline had a similar polypharmacological profile based on radioligand binding assays with the highest affinity for α1 (Ki=6.6 nM) and 5-HT_2B_ receptors (Ki=6.2 nM). Interestingly, N-desmethylxanomeline had potent interactions with other molecular targets including the 5-HT_1D_ (Ki=38 nM) and 5-HT_7_ (Ki=40 nM) serotonin receptors, various muscarinic receptors (Ki’s range from 93-219 nM) and many other GPCRs (Table 1). Taken together these results indicate that both xanomeline and its metabolite N-desmethylxanomeline have complex polypharmacological profiles with potent actions at many muscarinic, serotoninergic, dopaminergic, α1 and adrenergic receptors.

### Profiling xanomeline and N-desmethylxanomeline at 320 GPCRs reveals a large number of potential molecular targets

Given the complex polypharmacology of xanomeline and N-desmethylxanomeline, we next profiled them at 321 GPCRs using our PRESTO-Tango resource which provides an unbiased estimate of functional activity at the majority of druggable GPCRs in the human genome ^25^. In initial assays we screened xanomeline and N-desmethylxanomeline at 10 μM with a cut-off of potential activity at 2-fold change over basal as previously detailed ^26^. As shown in Fig 2, both xanomeline and N-desmethylxanomeline had potential activities at muscarinic, serotonergic, histaminergic and other GPCRs.

**FIGURE 2.**
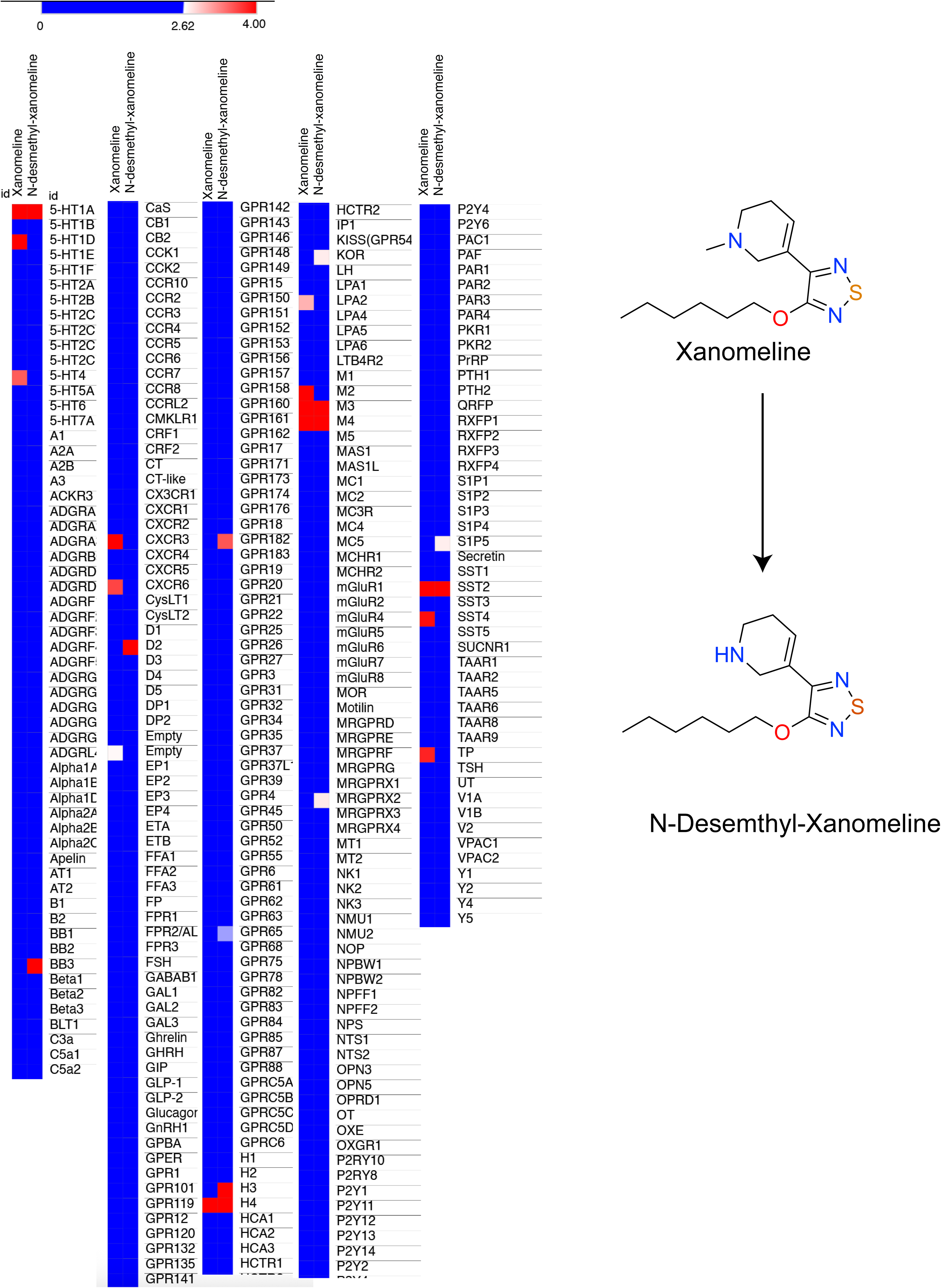
Results from PRESTO-TANGO screen of Xanomeline and N-desmethyl-Xanomeline. Shown in heat map format are the results from the initial PRESTO-TANGO screen at 10 μM final drug concentration. Data represent mean of 3 replicates. Shown to the left are the structures of Xanomeline and N-desmethyl-Xanomeline.

### Functional activity of xanomeline and N-desmethylxanomeline at muscarinic receptors

Given the robust potential polypharmacology of xanomeline and N-desmethylxanomeline, we next investigated each family of receptors separately using a variety of functional assays. We first focused on muscarinic receptors as they have been implicated in the beneficial antipsychotic activity of xanomeline ^27^ ^3^. For initial studies we examined the ability of xanomeline and N-desmethylxanomeline to activate various transducers for M1-M5 muscarinic receptors ^15^. To assess muscarinic receptor activation of arrestin translocation we used our PRESTO-Tango resource ^10^. For quantification of G protein activation, we used our BRET-based resource TRUPATH ^28–30^ and the results are found in Fig 3 and Table 2.

**FIGURE 3.**
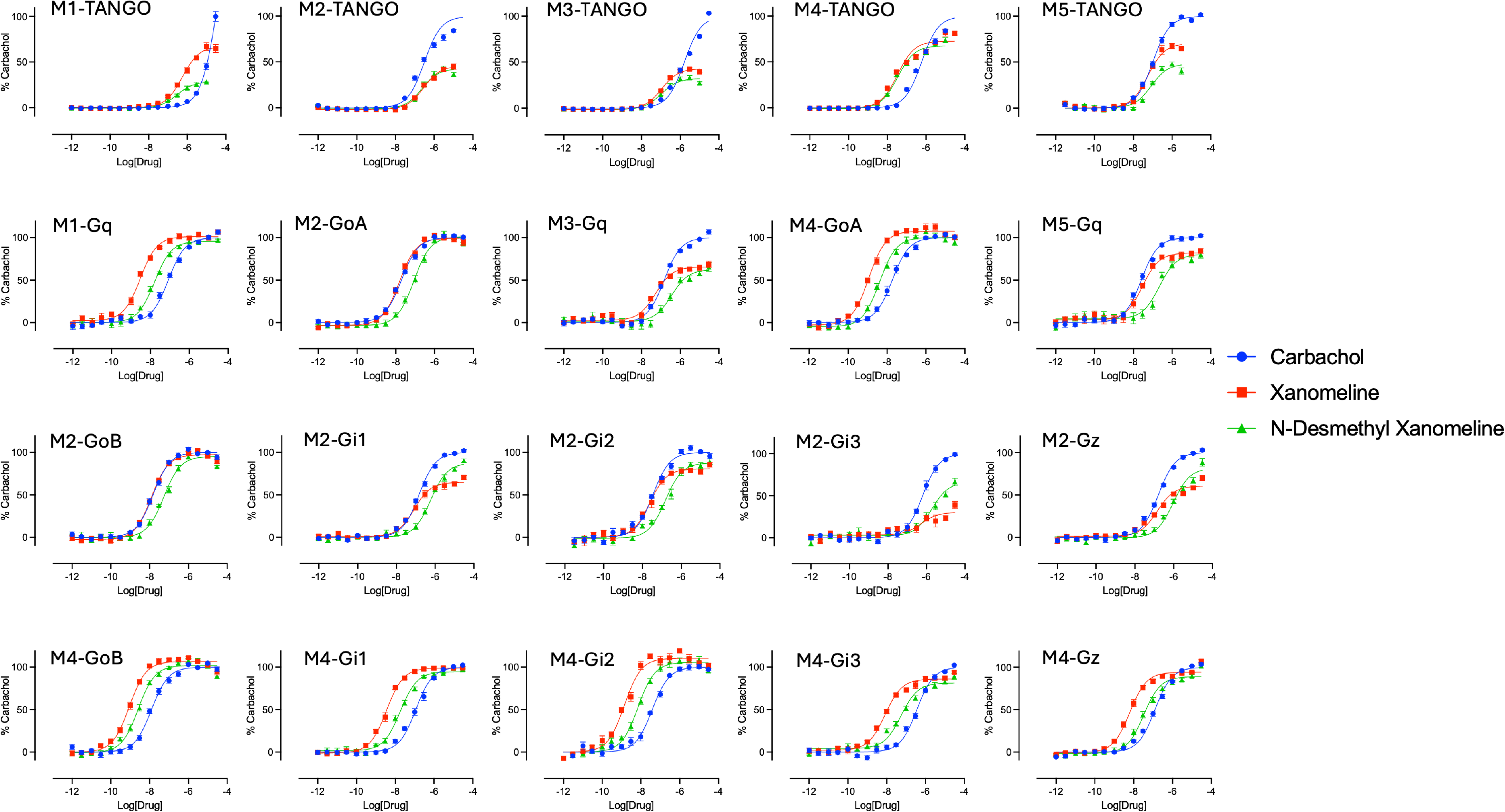
Concentration-response studies reveal differential activation of M1-M5 muscarinic receptor transducers. Shown are mean +/− S.D. of concentration-response studies, each of which incorporates results from 3 technical and 3 biological replicates, for carbacol (control muscarinic agonist), Xanomeline and N-desmethyl-Xanomeline using PRESTO-TANGO β-arrestin recruitment assays and TRUPATH G protein biosensors.

As can be seen (Fig 3), the relative efficacies and potencies of xanomeline, N-desmethylxanomeline and carbachol varied considerably across muscarinic receptors, with xanomeline being more potent and efficacious than carbachol at M1 and M4 muscarinic receptors, for example. Interestingly, N-desmethylxanomeline was less efficacious than xanomeline for inducing arrestin translocation at M1, M2, M4 and M5 receptors (Fig 3; Table 2).

### Functional activity of xanomeline and N-desmethylxanomeline at other molecular targets

We next evaluated the functional activity of xanomeline and N-desmethylxanomeline at additional molecular targets identified in the PRESTO-Tango screen from Fig 2 (Supplementary Table 1). For these studies, concentration-response studies were performed at several potential off-targets with results shown in Figs 4 and 5.

**FIGURE 4.**
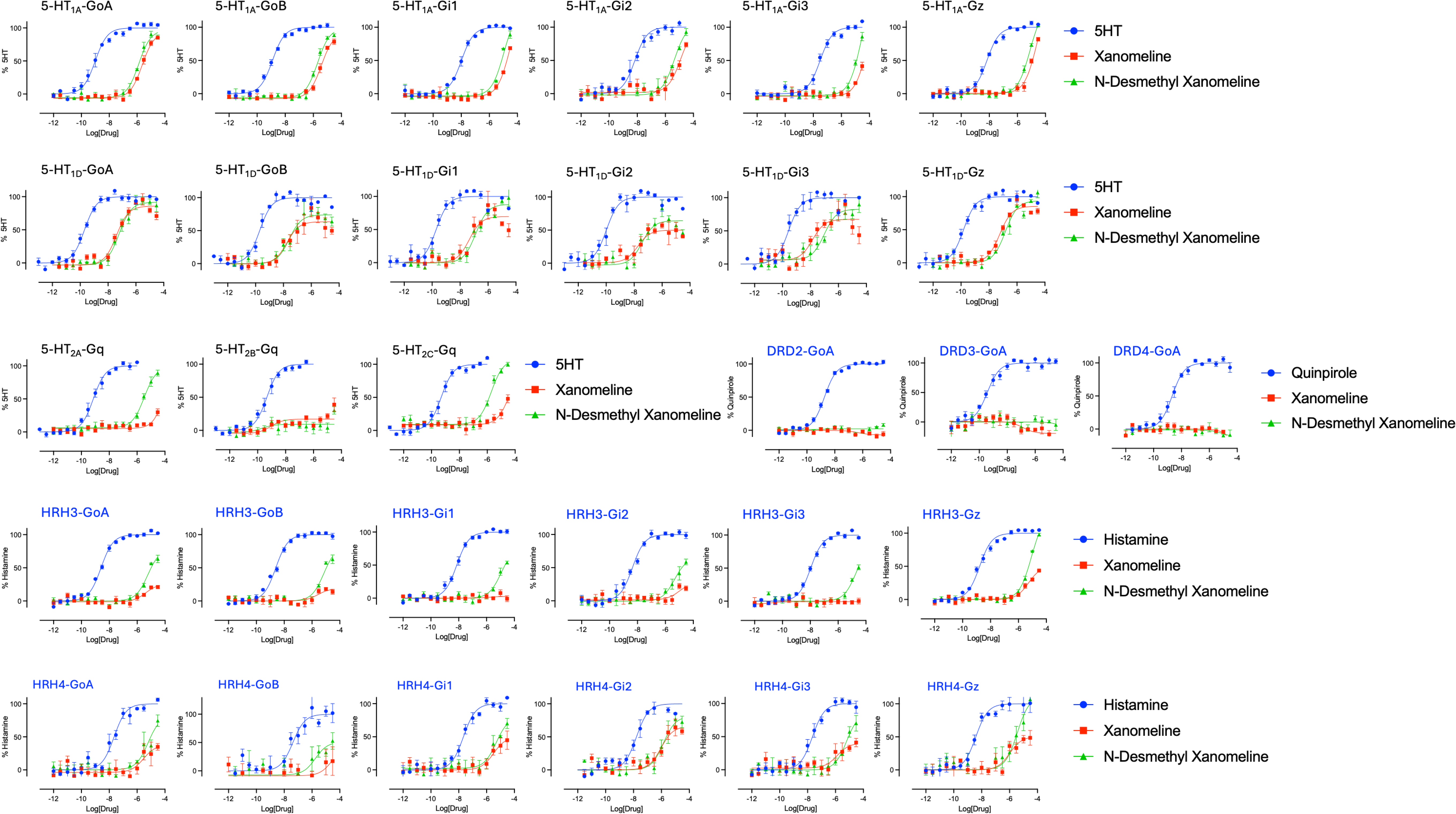
Concentrations-response studies reveal activities at serotonergic, dopaminergic and histaminergic receptors. Shown are mean +/− S.D. of concentration-response studies, each of which incorporates results from 3 technical and 3 biological replicates, for carbacol (control muscarinic agonist), Xanomeline and N-desmethyl-Xanomeline using TRUPATH G protein biosensors.

**FIGURE 5.**
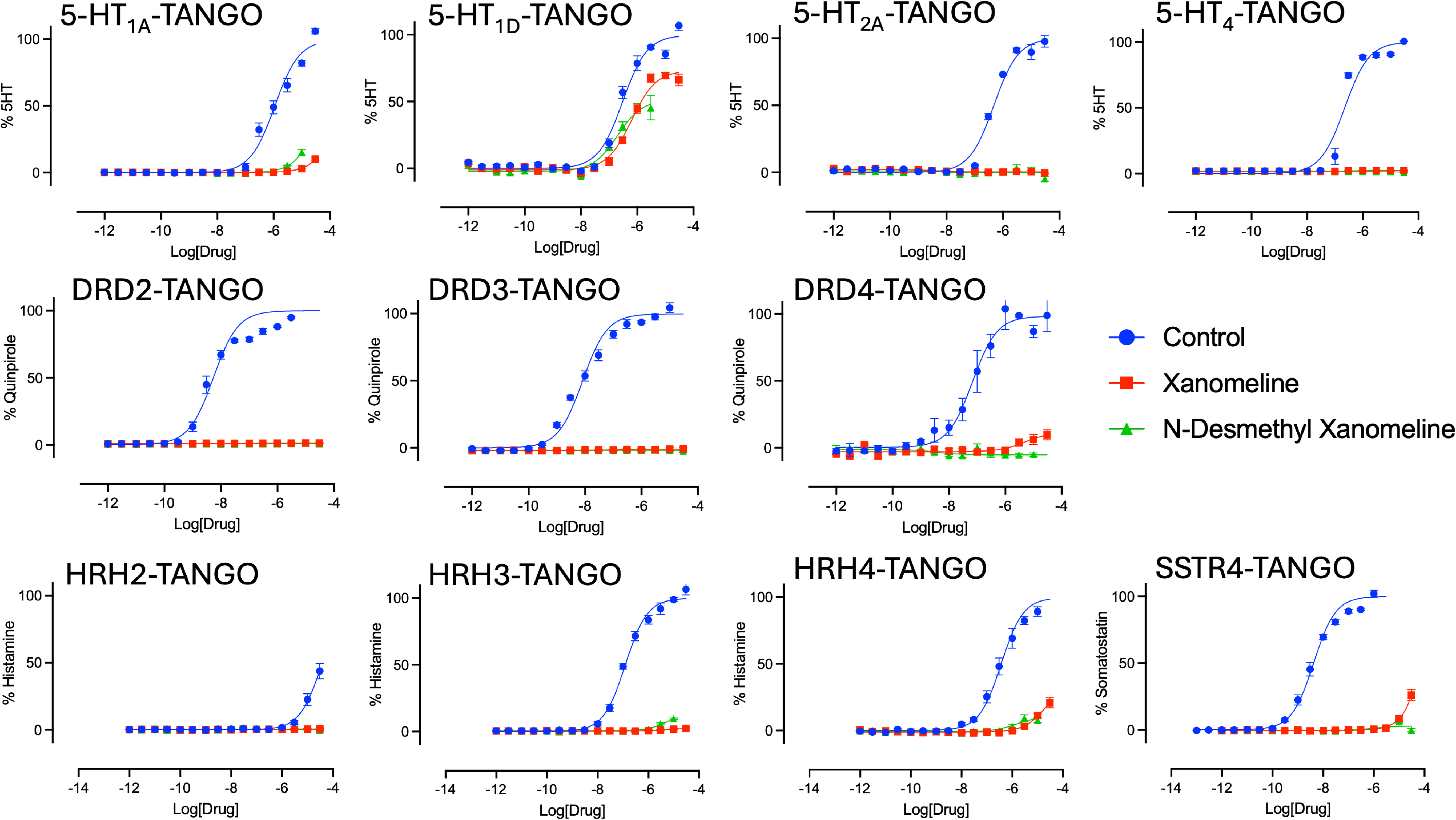
Concentration-response TANGO assays for Xanomeline and N-desmethyl-Xanomeline. Shown are mean +/− S.D. of concentration-response studies, each of which incorporates results from 3 technical and 3 biological replicates, for carbacol (control muscarinic agonist), Xanomeline and N-desmethyl-Xanomeline using PRESTO-TANGO β-arrestin recruitment assay

As can be seen, both xanomeline and N-desmethylxanomeline were full or partial agonists at 5-HT_1D_ and 5-HT_1A_ in a transducer-dependent fashion (Fig 4). The potencies for both compounds ranged from 17-130 nM with efficacies ranging from 48-91% (Fig 4 and Table 2) for 5-HT_1D_, while for 5-HT_1A_ potencies were much lower (EC50’s 2-20 μM). Xanomeline and N-desmethylxanomeline were inactive at other 5-HT receptors, although weak partial agonism for N-desmethylxanomeline was found for 5-HT_2A_- and 5-HT_2C_-serotonin receptors. Xanomeline and N-desmethylxanomeline had weak partial agonist activity at H3- and H4-histamine receptors with transducer-dependent potencies and efficacies.

We also evaluated other targets from the PRESTO-Tango screen and found no significant activity at the following targets: (HTR2A, HTR4, DRD2, DRD3, HRH2, GPR157, GPR182, and GPR25) with the results shown in Fig 5 and Supplementary Table 2. Finally, we calculated transduction coefficients ^31^ and clustered the values as shown in Fig 6.

**FIGURE 6.**
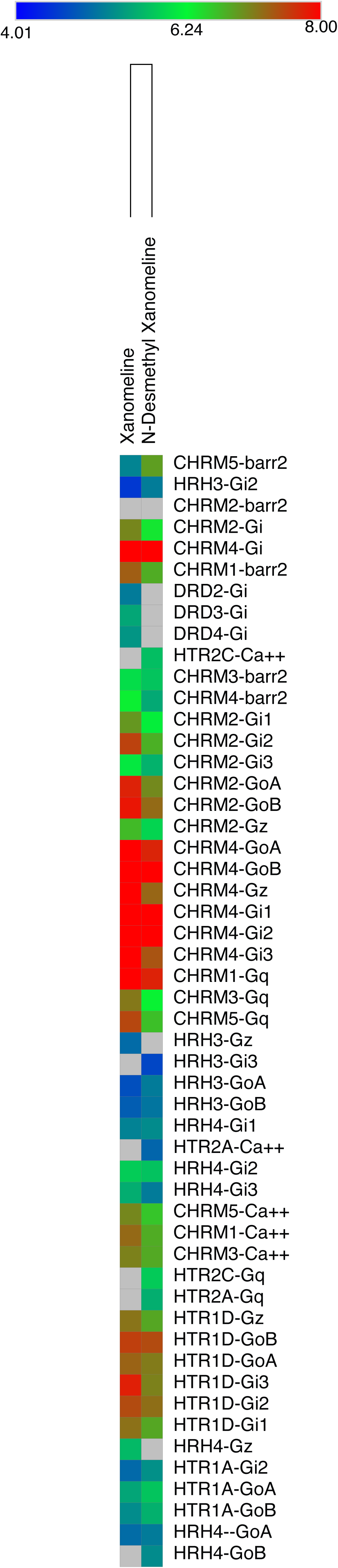
Heat-map of clustered transduction coefficients reveal multiple potential actions of Xanomeline and N-desmethyl-Xanomeline at muscarinic, serotonergic, histaminergic and dopaminergic receptors. Data are from Supplemental Table 1 with clustering performed with Morpheus. The values are color-coded according to the mapping at the top of the figure.

## Discussion

The main finding of this paper is that xanomeline and one of its metabolites N-desmethylxanomeline have complex polypharmacological profiles with activities not only at muscarinic acetylcholine receptors but also several other biogenic amine GPCRs. N-desmethylxanomeline had a similar profile as xanomeline with greater potency at M4-muscarinic receptors. As well, N-desemethylxanomeline had lower potencies and occasionally efficacies compared with xanomeline at other muscarinic receptors. As the M4 receptor has been implicated in the antipsychotic actions of muscarinic agonists, the results might predict a more favorable profile with N-desmethylxanomeline compared with its precursor xanomeline.

In prior studies, we have provided evidence in favor of the hypothesis that the polypharmacological profiles of antipsychotic and antidepressant medications ^7^, psychedelics ^19^, some anxiolytics ^32^, anorectic agents like fenfluramine ^33^, some anti-Parkinsonian medications and anti-migraine medications ^34^, 3,4-methylenedioxymethamphetamine ^35^ and many other approved medications and drugs of abuse ^36^ are involved in their therapeutic and/or side-effects. It is not too surprising, that xanomeline and it’s active metabolite N-desmethylxanomeline also have complex polypharmacological profiles.

Given the affinities and potencies of xanomeline and N-desmethylxanomeline for several serotonergic, dopaminergic, adrenergic and histaminergic and σ receptors it is conceivable that off-target actions at these sites contribute to the therapeutic or side-effects of xanomeline. Future studies examining more selective M1/M4 agonists will be needed to fully clarify the relevance of these activities for the observed actions of xanomeline.

